# Improved potential analysis for inadequate ecological data

**DOI:** 10.1101/2024.02.25.581934

**Authors:** Babak M. S. Arani, Egbert H. van Nes, Marten Scheffer

## Abstract

Potential analysis is used in many ecological studies to infer whether or not an ecosystem can have alternative stable states, to estimate the tipping points and, to assess the resilience of ecosystems. The main reason behind its frequent use is that such a frequency-based analysis is a minimalistic modelling strategy and therefore, is well-suited for limited ecological data with measurement errors. It has been used extensively in tree cover studies to discern alternative states of savannah and forest, the regime upon which savannah and forest coexist and, their corresponding resilience. Classical potential analysis may produce biased results due to inherent theoretical and practical limitations. This letter introduces a more sophisticated approach to address these shortcomings and enhance predictive capabilities regarding ecological dynamics, especially when working with insufficient data whose incorrect analysis can significantly impact management strategies.

## Introduction

Ecosystems may shift to an alternative stable state from where recovery might be difficult [1]. It is, therefore, an active research area in ecology to infer the ecosystem states from data and see whether or not alternative stable states exist. Although it is easy to examine this problem in theoretical models [2], it is challenging to rigorously conclude from limited field data whether an ecosystem has alternative stable states. A practical solution which is well-suited to inadequate, i.e., short and low resolution, ecological data was proposed by Scheffer and Carpenter [3]. Based on their approach one of the strongest hints from the field data is that the frequency distribution of ecosystems with multiple stable states is expected to be multimodal, with each mode representing an ecosystem equilibrium. This technique, i.e., a ‘*frequency-based distribution analysis*’, has become prevalent in ecosystem studies, inspiring the widespread use of ‘*potential analysis*’ [4], a form of frequency distribution analysis, in ecological research and beyond [5-10].

Potential analysis is used frequently in tree-cover analysis using remotely sensed extensive spatial data [6,10,11] monitored at sparse time points over time. The main interest is to infer alternative states of savannah and forest across rainfall gradients, the values of tipping points, predicting the Maxwell-point where both savannah and forest are equally resilient [11], and to estimate the resilience of savannas and forests. Indeed, the use of such frequency-based techniques have their own limitations, for instance it is assumed that the frequency distribution of data reflects the stationary distribution of the (yet unknown) system and that a simplest model of stochasticity (additive Gaussian white noise) [12] is assumed. Nonetheless, this paper focuses on scenarios where the basic assumptions of potential analysis hold, and more advanced methods of system reconstruction are impractical due to inadequate data [13-15]. Thus, potential analysis serves as a minimal stochastic modelling framework to elucidate key system features when high-quality data are lacking. However, classic potential analysis suffers from three major drawbacks: 1) it fails to reliably estimate deterministic forces, impacting the accuracy of system state estimations, particularly alternative stable states and repellors, 2) it inadequately estimates the regime of stochasticity, crucial for understanding ecological resilience, and 3) it does not directly incorporate data, relying solely on data distribution. Resilience measures and indicators (recovery rate or engineering resilience [16], Holling measures [17], exit time [18]) are all highly sensitive and fragile to poor estimation of system states especially stochastic regimes. To address these shortcomings, we propose an enhanced potential analysis approach that incorporates data, data distribution and, data distribution derivative, leading to more accurate estimations of deterministic forces and stochastic regimes.

The term ‘potential analysis’ comes from the ‘potential conditions’ [19] in which only some systems, including all one-dimensional systems, fulfil (higher dimensional systems which obey the potential conditions have a dynamics which somehow resembles the one-dimensional systems). This means that this technique is mainly applicable to univariate data and some multivariate data whose underlying dynamics is assumed to have a univariate nature. Therefore, here we restrain the discussions to unidimensional systems.

### Classical potential Analysis in a nutshell

Classic potential analysis is a kind of frequency distribution analysis where the functional form of ecosystem dynamics is mainly estimated by the data distribution. A classical paper where ecologists typically use for potential analysis is by Livina et al [4]. This analysis rests on two rather strong assumptions. The first one is that it is assumed that the data distribution is at equilibrium and does not change by time [20], i.e., the data distribution coincides with the so-called ‘*stationary probability distribution’* of the underlying system. Clearly, such an assumption requires that the data time span be long enough such that the probability distribution can be assumed to be stationary, which is usually not the case in short time-series ecological data. Exceptions may arise when analysing extensive spatial data, such as Moderate Resolution Imaging Spectroradiometer Satellite (MODIS) tree cover data, using the space-for-time substitution technique [21], where this assumption may hold merit. The second assumption underlying potential analysis posits that statistical fluctuations can be adequately described by the simplest model, namely ‘*additive Gaussian white noise*’. This entails that 1) the noise intensity remains independent of the system state (additive), 2) the noise is assumed to be drawn from a Gaussian distribution, and 3) the noise lacks temporal autocorrelation (white). These assumptions are necessary for establishing a one-to-one correspondence between the equilibria of the deterministic system (i.e., the states where the system settles in the absence of perturbations) and the modes of the stationary probability distribution [12] (see Figure 1).

**Figure 1.**
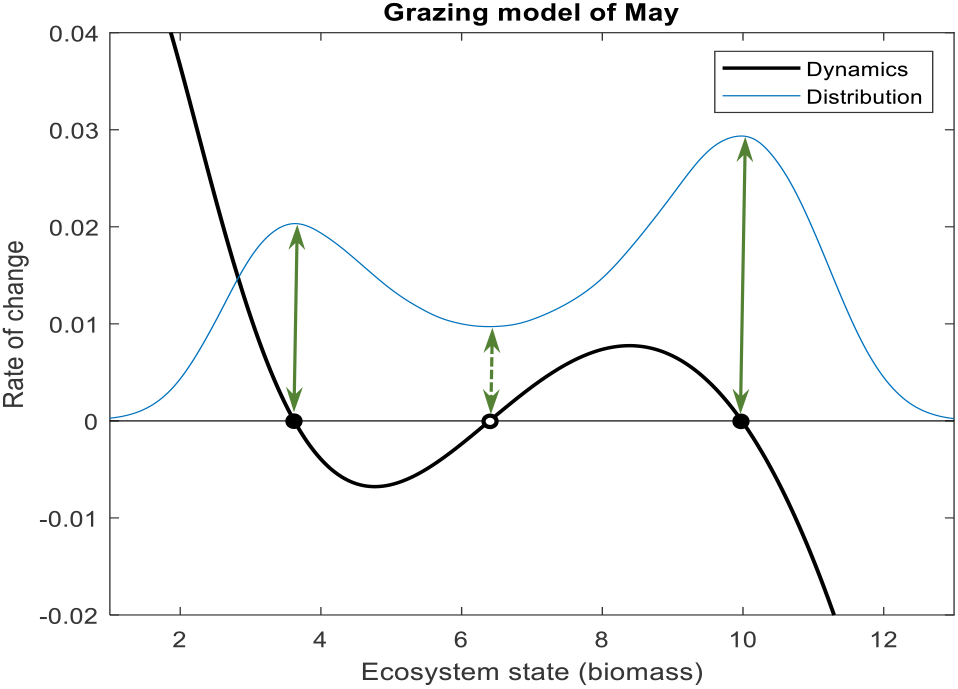
An illustrative example demonstrating the idea of classical potential analysis. Under this methodology the data distribution modes correspond with the equilibria and the distribution minima correspond with repellors in between. The black curve illustrates the rate of change, i.e., the dynamics, and the corresponding stationary distribution is illustrated in blue. Model is the grazing model of May [2], 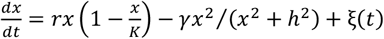 and the parameter values are r = 0.*1, K* = *2*0, r = 0.55r, *h* = r.r. r*(t)* is white noise with intensity of *σ* = 0.*1*r. The distribution is multiplied by 1/6 for a nicer representation.

Examples where these assumptions are violated includes: stochastic demography [22,23] and environmental stochasticity [22,24] where noise intensity depends on the population density (multiplicative noise), when extreme events happen to be more frequent as is the case with many natural disturbances like fires, floods, earthquakes, etc., in which fluctuations tend to have a fat-tailed distribution [25] (non-Gaussian noise), or when we oversimplify ecosystem states by not incorporating some other relevant state variables and this leads to coloured noise [26]. Detailed discussions about the effect of ‘*complex noise*’, i.e., when at least one of the assumptions 1-3 is violated, on the stationary distributions are outlined and discussed in detail in [12]. Here, unlike classical potential analysis, we relax assumption 1: we do not assume that the noise is additive (i.e., multiplicative noise. Note that when considering an additive noise model, the stochastic component is essentially the underlying time scale of the system). A consequence of this complex noise is that the mentioned one-to-one correspondence between the equilibria of the deterministic settings and the modes of stationary distribution might not hold. The core of our approach, however, will not change under either case of additive or multiplicative noises. Therefore, we give a rather detailed description for the case of additive noise and only show the corresponding results in case noise is multiplicative (but the full description is outlined in the appendix).

The idea of potential analysis consists of two steps: 1) Seeking a functional form for the deterministic component of the underlying system using the data distribution and an assumed functional form for the stochastic component 2) Estimating the regime of stochasticity (i.e., the noise parameters). Below, we provide a brief overview of both steps.

### Step 1: Finding a functional form for the system via data distribution

Assume that the following univariate stochastic model (called ‘Langevin’ or ‘Diffusion’ model in the literature) can describe the ecosystem dynamics:

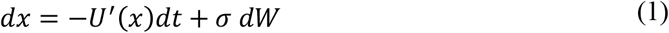

Where *x* is the ecosystem state (e.g., biomass) and *W* stands for the Wiener process (Thus, the noise source, *dW*, is white (uncorrelated) and has a standard Gaussian distribution) and, *U*^*′*^*(x)* is the derivative of the potential function *U(x)* with respect to *x. σ* is the noise intensity (or noise level). In (1) we have an additive noise since the noise intensity is a constant and does not depend on the state *x*. For a multiplicative noise the noise intensity is a function of state, i.e., *σ* = *σ(x)*. The potential function *U(x)* and the stationary probability density function *p(x)* meet the following relation [4,19] (see Appendix A)

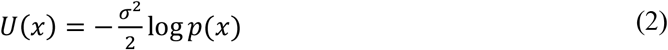

Formula (2) forms the core of the first step as it directly links the potential function *U(x)* (and therefore the system dynamics in (1) which requires *U*^*′*^*(x)*) to the stationary distribution *p(x)* which is approximated by the data distribution *p*_*d*_*(x)*. Therefore, (2) enables us to derive a functional form for the deterministic part of our system in (1) directly from the data distribution. The minima and maxima of *U(x)* correspond with the stable and unstable equilibria of (1) respectively. However, when we plug in the estimated data distribution *p*_*d*_*(x)* into (2) (i.e., the so-called ‘plug-in’ approach) to directly estimate *U(x)* by *p*_*d*_*(x)*, we may encounter numerous spurious local minima and maxima that do not align with true stable and unstable equilibria. This issue primarily arises due to two factors: 1) We have a finite sample of data, whereby in theory we need an infinite amount of data and 2) the presence of measurement errors in our data. Classical potential analysis relies on the mentioned plug-in approach. To more accurately estimate the true maxima and minima of *U(x)* and avoid overcounting the number of attractors, we need to reliably estimate the derivative of *U(x)* who’s zero crossings correspond to stable and unstable system states. In other words, we should reliably find the zero crossings of the slope of the negated potential function, i.e., *−U*^*′*^*(x)* in below, referred to as the ‘*ecosystem dynamics*’

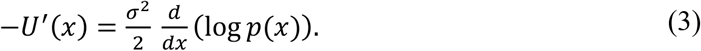

For the case of multiplicative noise there is a formula similar to (3) as bellow (see Appendix B)

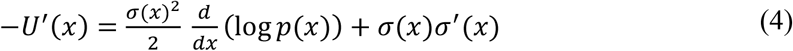

Where *σ*^*′*^*(x)* is the derivative of *σ(x)* with respect to *x*. So, in both additive and multiplicative cases we are left to estimate the derivative of data log-density reliably (this clarifies why we primarily focused on additive noise). The classical approach plugs in the data distribution *p*_*d*_*(x)* into (3) (or (4) in case of multiplicative noise) in this first step which may result in some spurious attractors. By the help of ecosystem dynamics in (3) we obtain the following functional form for our model in (1)

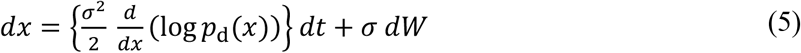

In the next section we adopt a reliable technique in estimating the derivative of data log-density which avoids plug-in methods.

### Step 2: Estimating the regime of stochasticity via data autocorrelation

The analysis of the first step leads us to a Langevin system (5). What remains to be estimated is a single noise parameter *σ*, i.e., the noise intensity. This parameter can be interpreted as the ‘time scale’ or ‘speed’ of (5). To this end, the classical potential analysis seeks to estimate *σ* via the autocorrelation function of data. However, other relevant such as ‘one-step conditional probabilities’ (for further details, refer to the appendix of reference [18], page 5) can serve the same purpose. In order to estimate the time scale of (5) one varies *σ* and compares the corresponding autocorrelation function of (5) with that of data. An optimal estimate of *σ* is obtained by finding the ‘best match’ (depending on the distance measure being used) between the two autocorrelations. This is indeed, an optimization problem. Using the autocorrelation function (or any other quantity that does not directly involve the data) to estimate *σ* can pose challenges. Firstly, the ‘best match’ lacks a unique interpretation and depends on the chosen metric for measuring the ‘distance’ between the model and data autocorrelations. Consequently, different distance measures may yield relatively different estimates for the optimal noise intensity. Secondly, while Langevin systems (1) typically follow an exponentially decaying autocorrelation function, the structure of the data autocorrelation function can be complex, making comparison difficult. Lastly, the final estimate is highly sensitive to the number of autocorrelation lags one choses to include in the process of comparing the autocorrelation functions of data and model. These challenges can significantly impact the accuracy of the estimated noise level, thereby affecting our ability to predict the system behaviour, particularly the resilience of the system against perturbations. In the next section we present an elegant approach to estimating the noise intensity for both additive and multiplicative cases by directly incorporating data.

## An advanced potential Analysis

As we observed in the previous section the classical potential analysis has caveats in both steps: it relies on a simplistic plug-in approach to derive a functional form for the system in the first step, and it estimates the system’s time scale using quantities such as autocorrelation of data, instead of involving the data directly. Here, we propose fundamental improvements to both steps.

### Step 1: Finding a reliable functional form for the system via data distribution and its derivative

As we saw in (3) or (4) we need to estimate the log-density of the stationary distribution, i.e., the quantity 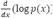, in both cases of additive and multiplicative modelling. A naïve way to do so is to plug-in the data distribution *p*_*d*_ into the log-density in (3) or (4) to arrive at (5). However, this approach may result in significant miscalculations (see Figure 2). To illustrate this more clearly, note that the log-density in (3) or (4) can be expressed as

**Figure 2.**
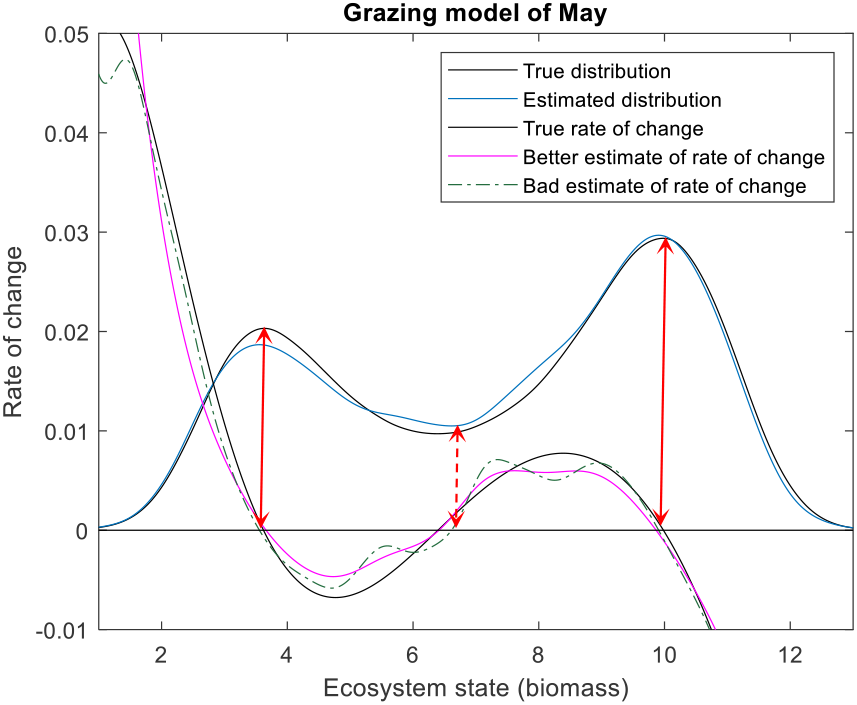
An example illustrating how the first step of the improved potential analysis works and how it differs from the classical approach. Using a sample of size 100000, the data distribution (blue) is estimated first. Following the classical approach, we obtain a non-reliable estimate for the rate of change, as the log density derivative (dot-dashed green) is not estimated reliably. However, by employing the improved analysis, we achieve a better estimate (purple). For clarity, we have illustrated the rate of change (i.e., *σ*^*2*^/*2* times the log density derivative, with *σ* = 0.*1*r being the true noise level) rather than the log density derivative itself. The red lines indicate the positions of alternate states and the repellor if we follow the classical approach. The data, model and its parameters are the same as in Figure 1.

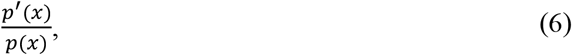

where *p*^*′*^*(x)* is the derivative of the probability distribution with respect to *x*. Therefore, inserting the estimated data distribution *p*_*d*_*(x)* into the log-density in (3) or (4) is equivalent to performing the same with (6), which involves dividing the derivative of the estimated data distribution by the estimated data distribution, i.e., 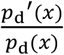. It is important to realize that ‘*estimation of the data distribution derivative does not equal the derivative of the estimated data distribution*’ [27] as we show later. Therefore, the estimation of log-density should not only involve the estimation of frequency distribution of data but also the estimation of the derivative of the distribution. This crucial fact is overlooked in classical potential analysis as it only estimates data distribution.

There are several methods to estimate distributions from sample data, among which kernel methods [28,29] are the most commonly used. Note that although we explain the idea of this section using kernel methods, the concept is independent of the specific technique used to estimate distributions. The kernel estimator of the unknown distribution *p(x)* is

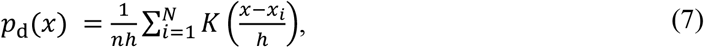

where the function *K* is called the kernel, *x*_*i*_ are the data points, *N* is the number of data and, *h* is called bandwidth which controls the smoothness of the estimator. A small bandwidth leads to under smoothing while a large bandwidth leads to oversmoothing. Therefore, the most crucial part of kernel methods concerns the correct estimation of bandwidth and there are several statistical approaches (for an overview see [30]) which find their usefulness in different applications. Here, to maintain consistency with the classical paper on potential analysis in [4], we consider the Silverman’s rule of thumb approach [31] where a Gaussian kernel *K* is used and the ‘*optimal bandwidth*’ *h*_*opt*_ is

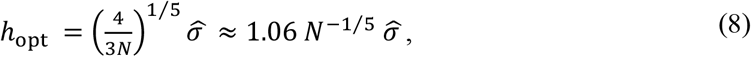

Where 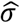 is the standard deviation of data. As we discussed earlier, we need an estimator for the derivative of data distribution in (6). An estimator *p*_*d*^′^_*(x)* for the data distribution derivative can be constructed by simply taking the derivative of *p*_*d*_*(x)* in (7), but the optimal bandwidth should be adapted acordingly. The ‘generalized’ Silverman’s rule of thumb (see [30,32] and its generalization to multivariate data and higher order derivatives of distributions, see also Appendix G) suggests the following optimal bandwidth *h*^*′*^_*opt*_ for the derivative of the data distribution

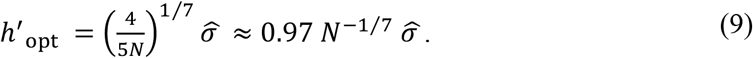

A comparison between *h*_opt_ and *h*^*′*^_opt_ in (8) and (9) reveals that *h*^*′*^_opt_ > *h*_opt_ for *N* ≥ 5 which is almost always the case. This significant result indicates that the estimation of the data density derivative requires a higher bandwidth, hence more smoothing, than that of the data density is needed. This implies that ‘*a correct estimation of log-density in (6) and therefore the ecosystem dynamics in (3) or (4), requires not only a correct estimation of the data distribution but also a correct estimation of the data distribution derivative*’. Figure 2 illustrates the performance of this approach using a simulated dataset from the grazing model of May with parameters being as in Figure 1.

One immediate important implication of (9) is that: ‘*A reliable estimate for the ecosystem states can be found by finding the zero crossings of the estimated data distribution derivative pd*^*′*^*(x), that is to say, one should solve for pd*^*′*^*(x)* = 0, *rather than finding the modes of the estimated data distribution p*_*d*_*(x)*’. Some other quantities of interest might require proper estimation of higher order derivatives of data distribution. For instance, the states where the system has the fastest recovery towards its equilibria (see the red stars in Figure 1, a measure of resilience, namely the minimum continuous input that is needed to bring the system to the alternative basin of attraction) require an estimation of the second derivative of data distribution as well. A similar analysis reveals that *h*^*′′*^_opt_ > *h*^*′*^_opt_ for *N* ≥ r where *h*^*′′*^_opt_ is the optimal bandwidth for the second derivative of the data distribution (in general, we need more smoothing as the order of density derivatives increases, see Appendix G).

Finally, in practice data are not independent as they exist as time series. Most methods for the estimation of distributions typically assume that data are independent. If our time series data is very large, then it is better to perform sparse sub-sampling first (depending on the strength of autocorrelations). Fortunately, correlations, even long-range ones, do not strongly affect the optimal bandwidth being calculated through techniques established for independent data [29], therefore we can safely use the distribution techniques developed for independent data.

### Step 2: Estimating the regime of stochasticity via data

Now that we have a reliable functional form for our system, we can proceed with estimating the noise level *σ* (if we wish to use an additive model) or, in general, the noise intensity *σ(x)* (if we wish to use a multiplicative model). In this step, unlike the classical potential analysis, we use the data directly. Here, we follow a maximum likelihood estimation (MLE) procedure, which is indeed an optimization problem (For more technical details we refer you to Appendix C&D, and here we only briefly describe the modelling details). MLE is a well-established inference technique once a model and its parameters are specified. This procedure can involve the entire data completely or a fraction of data if the dataset is very big and using the whole dataset only increases the computational time without significantly improving the final outcomes. An important property of MLE for Langevin models is that the uncertainty of the regime of stochasticity is lower compared with the deterministic part of the system [33-35]. Since the stochastic part of the potential analysis technique is also present in the deterministic part the use of maximum likelihood is very attractive.

When considering a multiplicative model, we consider two different modelling techniques to choose from. The first one involves considering a parametric form for the diffusion function *σ(x)*, for instance *σ(x)* = *a + bx*^*2*^, *a, b* > 0. This modelling strategy is useful when we have a ‘good’ justification to consider such a model for the regime of stochasticity. However, this is not often the case and we may have no clue about how *σ(x)* should look like in general. To address this problem, we have designed a second modelling technique which uses piecewise cubic polynomial forms called ‘cubic spline’ (for the details see Appendix E) under which cubic polynomial pieces are connected over a rather sparse and regular mesh of state space called ‘knots’ (see Figure 3, knots are the x-coordinate of red stars and the corresponding y-coordinates are the unknown parameters). The knot sequence can be irregular but we prefer a regular one as in this case the spline shows less sensitivity to the perturbations of the parameter values (i.e., the values of *σ(x)* at knots) [36] This is an elegant modelling technique splines are ‘flexible’ structures which help us to recover the unknown nonlinearities of *σ(x)*. Furthermore, splines have simple polynomial building blocks and this helps to solve the optimization problem in the MLE easier. Appendix E provides a detailed description of the superiority of the spline modelling technique. Figure 3A, illustrates the reconstruction of the stochastic regime for a simulated case study where we have used both modelling techniques.rFitting a multiplicative model to data may result in a mismatch between the stationary density modes with the equilibria of the deterministic part of the fitted model. This noise-induced phenomenon occurs due to the presence of multiplicative noise, which is a complex noise as elaborated on in the second section. For stochastic systems, the analogue of the ‘marble-in-a-cup’ stability landscape is the following ‘*effective potential*’ which provides a correct picture about the location of density modes

**Figure 3.**
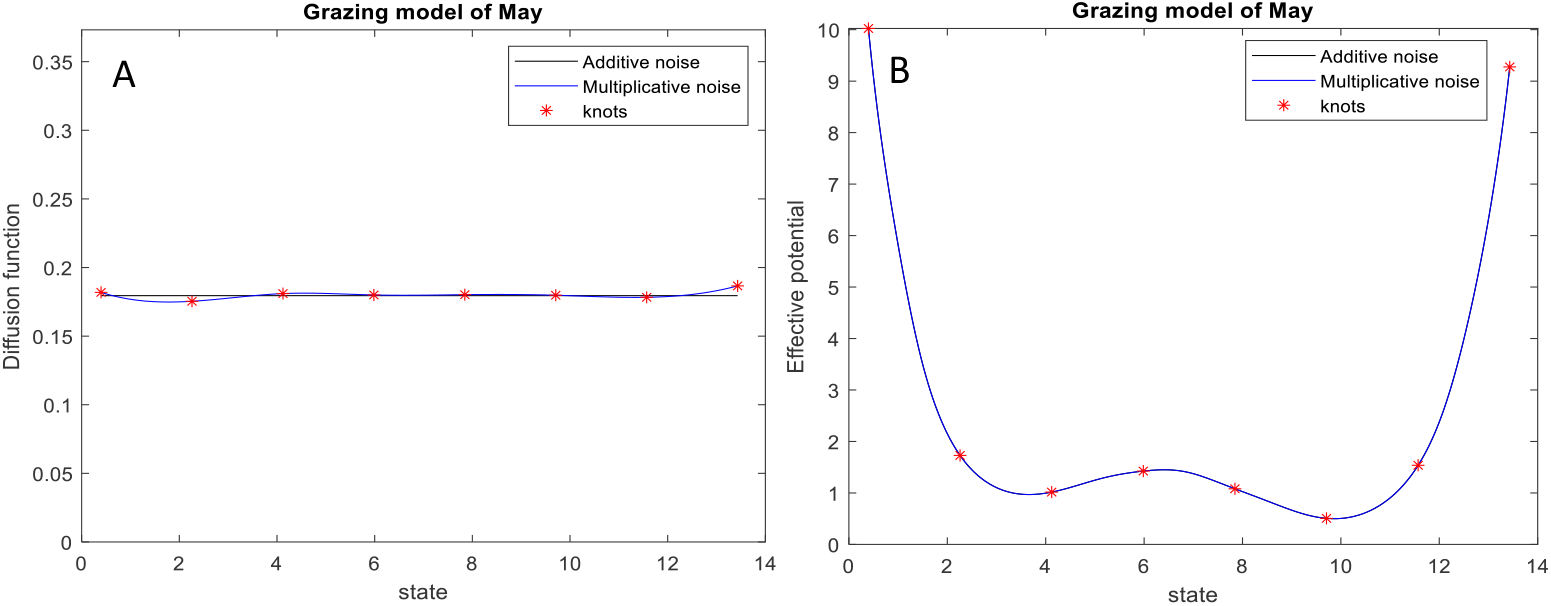
An example illustrating how the second step of the improved potential analysis works. (A) Here, we have fitted an additive model (black) as well as a multiplicative model using cubic splines (blue) via 8 evenly spaced knots (i.e., the x-values of the red stars) across the state space. Since in this simulated dataset we knew beforehand that noise is additive you do not see a big difference between the black and blue curves (otherwise, the differences could be higher). In the case of additive modelling the noise level is estimated as *σ* r 0.*1*r5 (the true noise level is 0.18) and its uncertainty is estimated as 0.0004. In the case of multiplicative modelling the estimation of *σ(x)* at 8 knots is [0.1819,0.1754,0.1810,0.1800,0.1802,0.1798,0.1783,0.1866]. Knots are connected with each other using cubic polynomial pieces. The corresponding vector of uncertainties is [0.0123,0.0015,0.0011,0.0012,0.0010,0.00086,,0.0013, 0.0117]. Note that the uncertainties of *σ(x)* at the first and last knots are the highest and this is expected since near the data borders there are less data points. (B) The corresponding effective potentials are shown where alternative states are the minima of the effective potential. The data, dynamic model and its parameters are as in Figure 1.

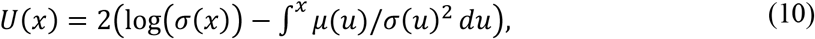

where the integral in (10) is an ‘indefinite integration’ and constant of integration is not important (i.e., what matters is the ‘potential difference’ between states rather than the corresponding values of potentials themselves). The effective potential is necessary to correctly estimate the location of alternative stable states, which are the potential minima and the repellors in between which are potential maxima (see Figure 3B).

### Tropical tree covers in South America and Africa: a case study

At intermediate range of precipitations (1000 to 2500 millimetres per year, see ([31])) forests and savannahs can be alternative biome states. The main reason is related to the positive feedbacks between fire and tree cover [31]: fire can maintain savannahs by supressing the establishment of forests [32] which in turn limits the fire spread as it requires a continuous grass layer [33]. Consequently, the distribution of tree cover is typically bimodal at intermediate range of precipitations (see Figure 4, left panel). However, existing tree cover datasets at continental scales are satellite data, which unfortunately are subject to measurement errors and exhibit rather high spatial autocorrelation, thereby reducing the effective number of independent data. This results in bumpy distributions although the overall patterns of tree cover are clearly bimodal. An immediate consequence is that naive approaches to estimating biome distributions and dynamics cannot accurately predict the rates of change in tree cover and the position of alternative stable states of savannah and forest. More critically, the location of repellor in between alternate biome states is uncertain (see Figure 4). The significance of the repellor lies in its role in correctly estimating resilience, as it represents the largest perturbation needed to trigger a critical transition. Therefore, understanding its location is crucial from a management perspective.

**Figure 4.**
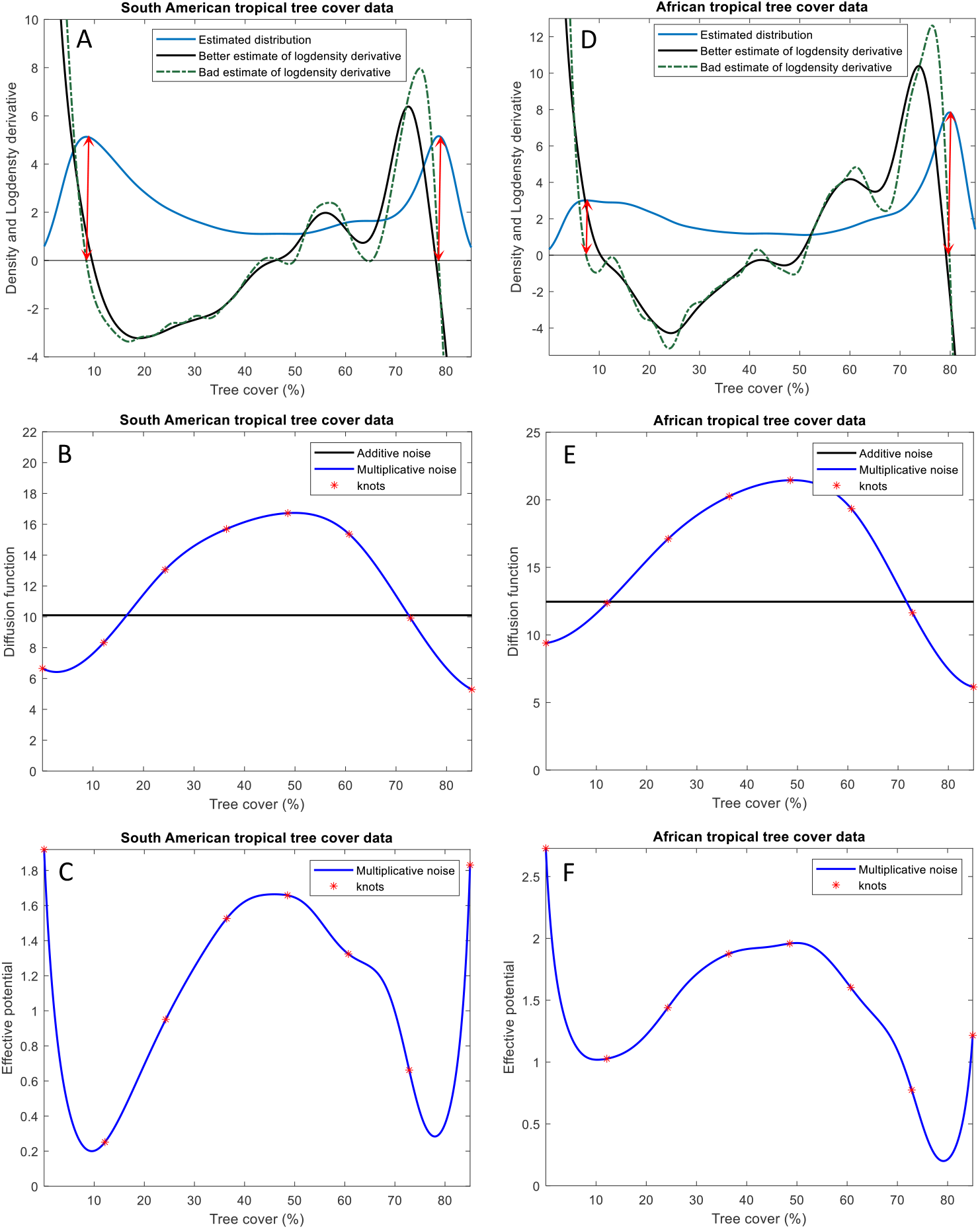
Application of the improved potential analysis to South American and African tropical tree cover data. (A&D) A non-reliable estimate of log density derivative (dot-dashed green) following classical approach versus a reliable one using improved potential analysis (black). The red lines show the positions of alternate states based on the classical approach. Due to non-reliable estimates following the classical approach there are many spurious alternate states and repellors. The density is multiplied by 200 for a nicer illustration (B&E) An additive stochastic regime (black) and a multiplicative one (blue) are fitted. Here, we have considered the time step of *dt* = *1* (arbitrary). In the case of additive modelling the noise level for South American and African data are estimated as *σ* r *1*0.*1*0 and *σ* r *12*.r5 with uncertainties estimated as 0.0032 and 0.0033 respectively. In the case of multiplicative modelling *σ(x)* is estimated to be [6.66 8.33 13.05 15.67 16.72 15.36 9.92 5.28] and [9.39 12.35 17.11 20.25 21.44 19.33 11.63 6.15] at 8 equidistant knots with the corresponding vector of uncertainties being estimated as [0.009 0.004 0.01 0.01 0.018 0.013 0.006 0.01] for south America and Africa, respectively. (C) The corresponding effective potentials with respect to the multiplicative modelling (for additive modelling the results are almost identical).

Here, we have analysed the South American and African tree cover data from Moderate Resolution Imaging Spectroradiometer Satellite (MODIS) with a 16-year annual resolution spanning from 2000 to 2016, at mean annual precipitation in the range 1639-1640 millimetres per year, respectively. We employed ‘space-for-time’ substitution approach [18] and made the assumption that all data within this short range of precipitation were subject to the same precipitation conditions. This assumption was necessary due to the limited availability of data for specific spatial locations. By treating all data under the same ecological conditions (i.e., the same level of precipitation) the ‘same’, we were able to obtain a larger dataset. As shown in Figure 4, the classical potential analysis yielded an unreliable estimate for the location of the repellor. Particularly, the classical methods seem to under-smooth the ecosystem dynamics. Our ‘improved’ estimator, however, produced satisfactory results. Furthermore, the estimates of the alternative stable states of savannah and forest deviate a bit from the corresponding tree cover modes.

## Conclusion

Potential analysis is a fascinating, parsimonious, and convenient technique for reconstructing the underlying system from data, as its theoretical foundations rely on simple assumptions about the regime of stochasticity [12]. The classical approach aims to fit a dynamical model to data by utilizing the data distribution in the first step and data autocorrelation in the second step, without directly involving the data itself. One immediate consequence of this is that the classical technique cannot distinguish between two different datasets which have similar distributions and correlation information. In contrast, we have developed a methodology that incorporates both the data distribution and its derivative in the first step, while fully involving the data in the second step. This approach allows for a more comprehensive analysis and better discrimination between datasets with similar statistical properties.

The challenge associated with the improper use of potential analysis, using the classic approach, become particularly pronounced when dealing with insufficient data. This poses a significant obstacle given the often highly limited availability of ecological data. Improper estimations of ecosystem dynamics resulting from such limited data can lead to inaccurate predictions of critical ecological quantities, including alternative stable states, tipping points, Maxwell points, attraction basins, and ecosystem resilience. These inaccuracies have profound adverse effects on ecosystem management. While our approach addresses these problems by improving the accuracy of potential analysis, it is important to emphasise the ongoing necessity and urgency of obtaining sufficient ecological data of high quality. Access to such data will enable the utilization of more advanced techniques for system reconstruction [12-15], representing an essential next step research in ecological research.

## Appendix A: Classical potential analysis in a nutshell

Consider the following one-dimensional diffusion (also called Langevin) model

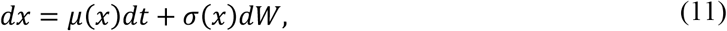

where *μ(x)* is referred to as the drift function and *σ(x)* as the diffusion function. *W* represents a standard Wiener process or Brownian motion, with ‘standard’ indicating that it has a zero mean and unit variance, resulting in the noise source *dW* being white (uncorrelated) and Gaussian. The diffusion function can be thought of as the ‘weight’ assigned to stochastic fluctuations, indicating the intensity of noise at a given state *x*. When the diffusion function does not vary with the state, the noise is termed ‘additive’, otherwise it is termed ‘multiplicative’. The classical potential analysis relies on additive noise, and we make this assumption here. Given a time series generated by this model, our goal is to estimate the drift and diffusion functions. Since (11) is one-dimensional it can always be rewritten as follows:

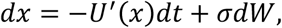

Where *U(x)* is called the ‘potential function’. The name ‘potential analysis technique’ is due to the presence of potential function, which is guaranteed to exist for one-dimensional systems. However, in higher dimensions, it often does not exist. The long-term distribution of (11), referred to as the stationary distribution [4,37], is given by

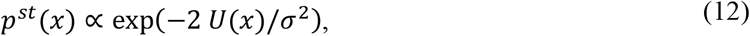

where ∝ denotes proportionality. Using (12), there exists a one-to-one correspondence between the stationary distribution and the potential function. A rather strong assumption of the potential analysis technique is that the dataset is sufficiently long to consider the data distribution *p*_*d*_*(x)*, as coinciding with the stationary distribution *p*^*st*^*(x)*, i.e., *p*^*st*^*(x)* = *p*_*d*_*(x)*. Consequently, we can solve (12) for the potential to obtain

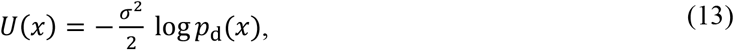

Equation (13) serves as a cornerstone of potential analysis technique, constituting the first step of this technique. It facilitates the reconstruction of the potential function *U(x)*, and consequently, the drift function of (11), *μ(x)* = *−U*^*′*^*(x)*, directly from the data distribution. The second step of potential analysis involves estimating the noise level. This is achieved through an optimization process where the optimal noise level *σ* is estimated by finding the ‘best’ match (depending on the considered distance measure) between the autocorrelation function of data and that of model (11).

## Appendix B: Improved potential analysis for multiplicative noise

Here, we relax one of the assumptions of potential analysis by allowing the noise to be multiplicative, meaning that the diffusion function can depend on the state *x*. The relation (12) can be generalized to the case of multiplicative noise as *[37]*

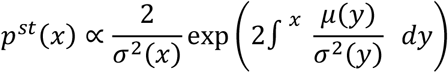

Where the integral is ‘indefinite integration’. We can solve the previous relation for *μ(x)* to get

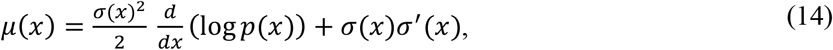

We use relation (14) as a basis for the first step of the ‘improved’ potential analysis.

## Appendix C: maximum likelihood estimation (MLE) of diffusion models

Consider a diffusion model

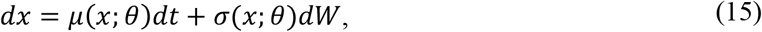

where *θ* is the vector of unknown parameters. Since we are following the potential analysis technique here, we are only concerned with estimating the diffusion function *σ(x*; *θ)* in (15), as the drift term *μ(x*; *θ)* can be obtained by the diffusion function and data distribution based on the relation (14). Assume that we have observations {*X*_0_, *X*_*1*_, …, *X*_*N*_} of this process at discrete times *t* = 0,*1*, …, *N* for a particular set of parameter values which we aim to estimate, and let Δ be the sampling time. Additionally, assume that *P*_*X*_(Δ, *x*|*x*_0_; *θ)* represents the conditional density of *X*_*t+*Δ_ = *x* given *X*_*t*_ = *x*_0_. The likelihood function for this set of observations is the multi-point density *P*_*X*_*(X*_0_, *X*_*1*_, …, *X*; *θ)*. Due to the Markovian nature of diffusion model (15) and using Bayes rule in probability theory, we can break this multi-point density by the products of the conditional densities as follows:

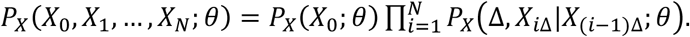

Based on the principle of maximum likelihood estimation (MLE), the true parameters of the diffusion model (15) correspond to the global maximum of the likelihood function. It is more convenient to work with the following log-likelihood function

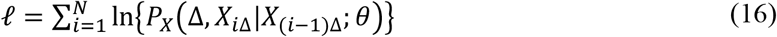

Where the first unconditional term In*(P*_*X*_*(X*_0_)) is ignored as it has a negligible effect when *N* ≫ *1*. However, we include it when the dataset at hand consists of multiple separate time series (i.e., replicate data), instead of a single series, each having its own initial observation *X*_0_ when the number of such series is rather high. In this work, we actually consider the negative log-likelihood function *−ℓ* and solve a minimization problem, instead.

## Appendix D (Univariate Euler reconstruction)

The Euler reconstruction corresponds to a maximum likelihood estimation of the diffusion model (15) when the sampling time is infinitesimally small, i.e., Δr 0. Under this limiting case we can replace our diffusion model with a difference equation based on the Euler-Maruyama discretization

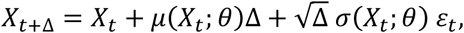

where *ε*_*t*_ follows a standard normal distribution. As a result, the conditional density *P*_*X*_(Δ, *x*|*x*_0_; *θ)* can be approximated by the following normal distribution with mean *x*_0_ *+ μ(x*_0_; *θ)*Δ and variance Δ*σ*^*2*^*(X*_0_; *θ*)

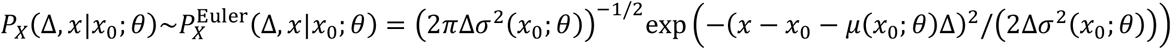

The big advantage of using the Euler approach is due to its low computational burden. Fortunately, based on our extensive experience, this approach also demonstrates satisfactory performance for datasets with medium resolution.

## Appendix E. Spline forms as an elegant implementation technique for univariate data with several reasons for their superiority

The inference technique we are employing is parametric, manning that one should specify functional forms for the diffusion function *σ(x*; *θ)* in (15) prior to embarking on the MLE. Note that here we are only concerned with the estimation of diffusion function in (15), as the drift function can be derived from the diffusion function and data using (14), as we adhere to the potential analysis technique.

While polynomials may initially seem appealing as candidate models for *σ(x*; *θ)*, they are not the best. High-order polynomials may be required to adequately represent complex nonlinearities in the diffusion function, resulting in a large number of parameters to estimate. Instead, piecewise polynomial forms, known as splines, offer a more suitable alternative. In more technical terms, a spline is a function composed of several polynomial pieces that are joined together at a sequence of points {*x*_*1*_, *x*_*2*_, …, *x*_*M*_} called a ‘*knot sequence*’. These polynomial pieces are arranged in such a way that the resulting function satisfies certain ‘smoothness’ conditions at knots. In spline modeling, the parameters consist of the unknown values of the diffusion function at the knots. Since in practice we do not have an extremely long dataset, and we prefer a model with a relatively low number of parameters, we consider a relatively sparse knot sequence across the state space, which typically corresponds to the data range. However, in situations where the number of data points near the data borders is very low, we consider a state space slightly smaller than the data range and treat those few data points falling outside this range as missing values. To distinguish between reconstruction procedures based on the typical parametric forms and those based on spline forms, we refer to the former as ‘*parametric reconstruction*’ and the later as ‘*spline reconstruction*’.

Here, we elaborate on several other reasons for the superiority of spline forms. The second reason is that, in general we often lack a proper understanding of a suitable parametric model form to try, unless there is strong empirical or theoretical justification. Even if one intends to pursue a parametric reconstruction, the flexibility of spline forms can provide valuable guidance in selecting appropriate candidate parametric models to try. The third reason is that during the process of estimating the model parameters using the MLE, the optimization algorithm is restricted to search within a relatively small parameter space containing the true but unknown parameters. In general, it can be challenging to identify an appropriate bounded parameter region that includes the true solution when using a parametric model. In contrast, spline models offer the advantage of easily defining a proper bounded region for the reconstruction algorithm to explore. This convenience arises because, in spline reconstruction, our parameters have a specific meaning to us, as they are the values of diffusion function at knots. Therefore, to estimate the diffusion parameters, we can define a search region like (0, *L*] for some *L* > 0 for all noise parameters, as the diffusion function should be positive. If we normalize the data beforehand, we can select a rather small value for *L*, such as *L* ≤ 5, and this means that the optimization algorithm can search within a rather small region. The fourth reason is that using splines ensures that our model does not exhibit global sensitivity to parameters as changes in a parameter value at a single knot do not propagate across the entire knot sequence. Instead, the effect is localized over an evenly spaced knot sequence, which we typically choose. The rationale for selecting a regular knot sequence, besides its convenience, is that a typical spline interpolation with (x-data, y-data) is guaranteed to have low sensitivity to perturbations of y-data (i.e., the model parameters) if the chosen knot sequence is equidistance (for details, see the reference [36]). This alleviates the pressure on the optimization algorithm, which in turn improves both the accuracy and speed of the optimization process, reducing the risk of failure. The last reason is that the spline function *σ(x*; *θ)* is surprisingly a linear function of parameters *θ* (although it is a nonlinear function of state *x*), and this property is quite helpful for estimating the parameters more easily and quickly.

## Appendix F. Gradient descent and grey wolf optimizer as local and global optimization algorithms used to solve the MLE

In order to estimate the parameters of the model (**Error! Reference source not found**.) using the MLE framework, we should find the global minimum of the negative log-likelihood function *−ℓ* where *ℓ* is defined in (16). Note that this is a constrained optimization problem since we need to ensure that *σ(x)* > 0. We used a simple ‘death penalty’ technique to handle this constraint. To tackle this global optimization problem, we resorted to meta-heuristic optimization algorithms. Specifically, we used the Grey Wolf Optimizer (GWO) [38], actually an improved version of it [39]. GWO is a novel meta-heuristic algorithm which is quite powerful in terms of both exploration (global search) and exploitation (local search). Fortunately, since splines have simple polynomial building blocks and are linear in the vector of parameters *θ*, the corresponding objective function, i.e., *−ℓ*, does not often have a complex structure and geometry. Because of this simplicity, we also used gradient descent algorithms, along with several starting points. Gradient descent algorithms are local solvers but are faster than GWO.

### Accessing the uncertainty of the parameters

To calculate the variance of the vector of estimated parameters, say 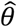, we need the Fisher information (FI) matrix given by

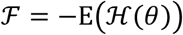

Where *H(θ)* = *∂*^*2*^*ℓ*/*∂θ∂θ*^*T*^ is the Hessian matrix, i.e., the second order partial derivatives of the log-likelihood function with respect to *θ*. In our minimization case, whewre we use *−ℓ* as our objective function, the FI matrix does not have a negative sign. The FI matrix is approximated by the ‘observed’ Fisher information as 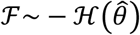. Then, the variance-covariance matrix *Σ* = *F*^*−1*^, i.e., the inverse of the Fisher information matrix, can be estimated as 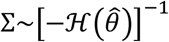. Eventually, the standard error for the i^th^ parameter, Say 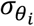, which is the square root of the i^th^ diagonal element in *Σ*, can be estimated by the square root of the i^th^ diagonal element of 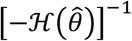.

## Appendix G Generalized Silverman’s rule of thumb for high dimensions and high order derivatives

Assuming *X*_*1*_, *X*_*2*_, …, *X*_*n*_ are an i.i.d (i.e., independently and identically distributed) sample from an unknown *d*-variate distribution, the multivariate kernel density estimation, as a generalization of univariate kernel density estimation, i.e., the relation (7) in the main text, is given by (see [30,32])

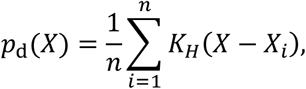

where *K*_*H*_*(X)* = |*H*|^*−1*/*2*^*K(H*^*−1*/*2*^*X)*, |. | means determinant, *H* is the *d*-by-*d* ‘bandwidth matrix’, and *K* is the multivariate kernel function. Typically, a Gaussian kernel 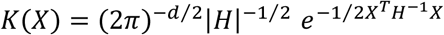 means transpose) is considered. Therefore, the bandwidth matrix *H* can be interpreted as a covariance matrix in this context. Generalizing Silverman’s rule of thumb for the r^th^ derivative of a *d*-variate distribution suggests the ‘optimal’ bandwidth matrix *H* to be diagonal with diagonal elements 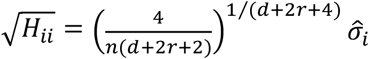 where 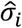 is the standard deviation of the i^th^ variable [32].r

